# Defining early human developmental identity: A curated, cross-platform marker framework

**DOI:** 10.64898/2026.03.26.714482

**Authors:** Jochen Dobner, Alessandro Prigione, Andrea Rossi

## Abstract

Early human development involves dynamic transitions in cell identity, including transient transcriptional modulation and stable lineage commitment. Distinguishing these types of gene expression changes is challenging and can be further exacerbated by genetic and experimental heterogeneity in the context of human pluripotent stem cell (hPSC) research. To address this challenge and help uncover transcriptional changes indicative of true developmental state, we establish a curated, cross-platform marker framework for robust identification of pluripotency and early germ-layer identity. Starting from an unbiased RNA-seq discovery set, we systematically validate candidate markers across qPCR, bulk and single-cell RNA sequencing, and quantitative proteomics platforms, yielding a refined panel of 67 markers (20 for the undifferentiated state, 17 for endoderm, 15 for ectoderm, and 15 for mesoderm). We show that this framework reliably identifies early developmental states across heterogeneous datasets, generalizes to in vivo human embryo cell types, and preserves lineage identity despite substantial transcriptional variability. Furthermore, we demonstrate concordant protein-level expression for a subset of markers, supported by deep proteomic profiling of the reference line KOLF2.1J. To enable broad application, we introduce DeepDiff, a web-based resource integrating the validated markers, allowing automated fate classification in a user-friendly interface. Together, this work provides a standardized framework for defining early human developmental identity and disentangling lineage commitment from context-dependent modulation.

## Introduction

Early human development is defined by the progressive establishment of cell identity, as pluripotent cells traverse transient and partially overlapping states toward lineage commitment (Boroviak et al., 2015; Rossi et al., 2016). These transitions are continuous and context-dependent, complicating the assignment of discrete cell identities. Experimental systems that model these processes therefore rely on molecular markers to infer cell identity, developmental progression, and responses to perturbation (Bock et al., 2011; Dobner et al., 2024a; Schmidt et al., 2023; Tsankov et al., 2015).

Human induced pluripotent stem cells (hiPSCs) have become a central platform for studying early development and disease (Takahashi et al., 2007). Their ability to differentiate into virtually all somatic lineages makes them highly relevant for investigating developmental processes and therapeutic interventions (Kirkeby et al., 2025). This is reflected by a rapid expansion of the field with over 16,000 publications reported between 2019 and 2024. However, in vitro models based on hiPSCs introduce variability arising from differentiation protocols, genetic backgrounds, and environmental conditions. Consequently, transcriptional states can shift without reflecting true lineage commitment, complicating the interpretation of experimental outcomes. While initiatives such as the International Society for Stem Cell Research (ISSCR) provide marker recommendations for pluripotency and trilineage assessment, these markers have not been systematically curated or validated across experimental platforms and directed differentiation contexts (Allison et al., 2018; Bock et al., 2011). This limits comparability across studies and hinders the reliable assessment of cell identity (Kamimoto et al., 2023; Morris, 2019; Tung et al., 2017).

Building on our ongoing efforts to define robust markers of early human development, we previously applied long-read RNA sequencing to identify transcriptional signatures associated with early germ-layer identity, resulting in 172 candidate marker genes not covered by the ISSCR guidelines (Dobner et al., 2024a). A subset of these markers was validated by qPCR and integrated into the hiPSCore framework, demonstrating the feasibility of reproducible and standardized classification. However, this initial framework was not designed to capture subtle variation within lineages or to assess marker performance across experimental contexts. As a result, it remained unclear which markers reflect stable lineage identity and which are influenced by experimental variation, such as environmental stressors or pollutants.

Here, we systematically curate and validate a cross-platform marker framework for early human developmental identity. Starting from an RNA sequencing–based discovery set, we filtered and validated candidate markers across qPCR, bulk and single-cell RNA sequencing, and quantitative proteomics, yielding a refined panel of 67 genes with consistent performance across platforms and experimental contexts. This curated marker set robustly identifies early developmental states across heterogeneous datasets, captures intra–germ-layer heterogeneity, and generalizes to single-cell transcriptomic data from human embryo development. Notably, lineage identity remains detectable despite substantial transcriptional variability, underscoring the need to distinguish stable identity from context-dependent modulation. To enable broad application, we provide DeepDiff, an accessible web-based resource for interrogating gene expression and protein dynamics across differentiation states. Together, this work establishes a standardized and transferable framework for defining early human developmental identity and its modulation by genetic, environmental, and experimental perturbations.

## Results

### qPCR-based validation of candidate marker genes for early developmental states

To define reliable marker genes for early developmental states, we performed independent validation of candidate genes identified in our previous RNA sequencing–based discovery effort (Dobner et al., 2024a). Using four to six, commercially available hiPSC lines per target gene, we systematically assessed marker performance by qPCR across undifferentiated (n = 18), endoderm (n = 17), ectoderm (n = 15), and mesoderm (n = 18) differentiated samples.

From an initial set of 172 candidates, we selected 68 genes for validation (Supplementary Table 1). This subset was designed to ensure robust representation across all differentiation states and to include a sufficiently large panel of markers per lineage (≥10 genes) to account for potential context-dependent variability, including genetic perturbations such as gene knockouts that may compromise individual marker reliability as previously described (Fischer and Gillis, 2021).

Marker performance was evaluated based on the ability to distinguish target from non-target cell states. Across all lineages, the majority of candidate markers showed significant enrichment in their respective target states (Fig. 1, Supplementary Table 2). All undifferentiated markers (17/18) were validated, alongside 82% (14/17) of endoderm, 80% (12/15) of ectoderm, and 67% (12/18) of mesoderm markers (p ≤ 0.05). Notably, one undifferentiated marker, *HHLA1*, despite being significantly differentially expressed, exhibited opposite regulation in qPCR compared to RNA-seq (downregulated vs. upregulated) and was therefore classified as not validated.

**Figure 1.**
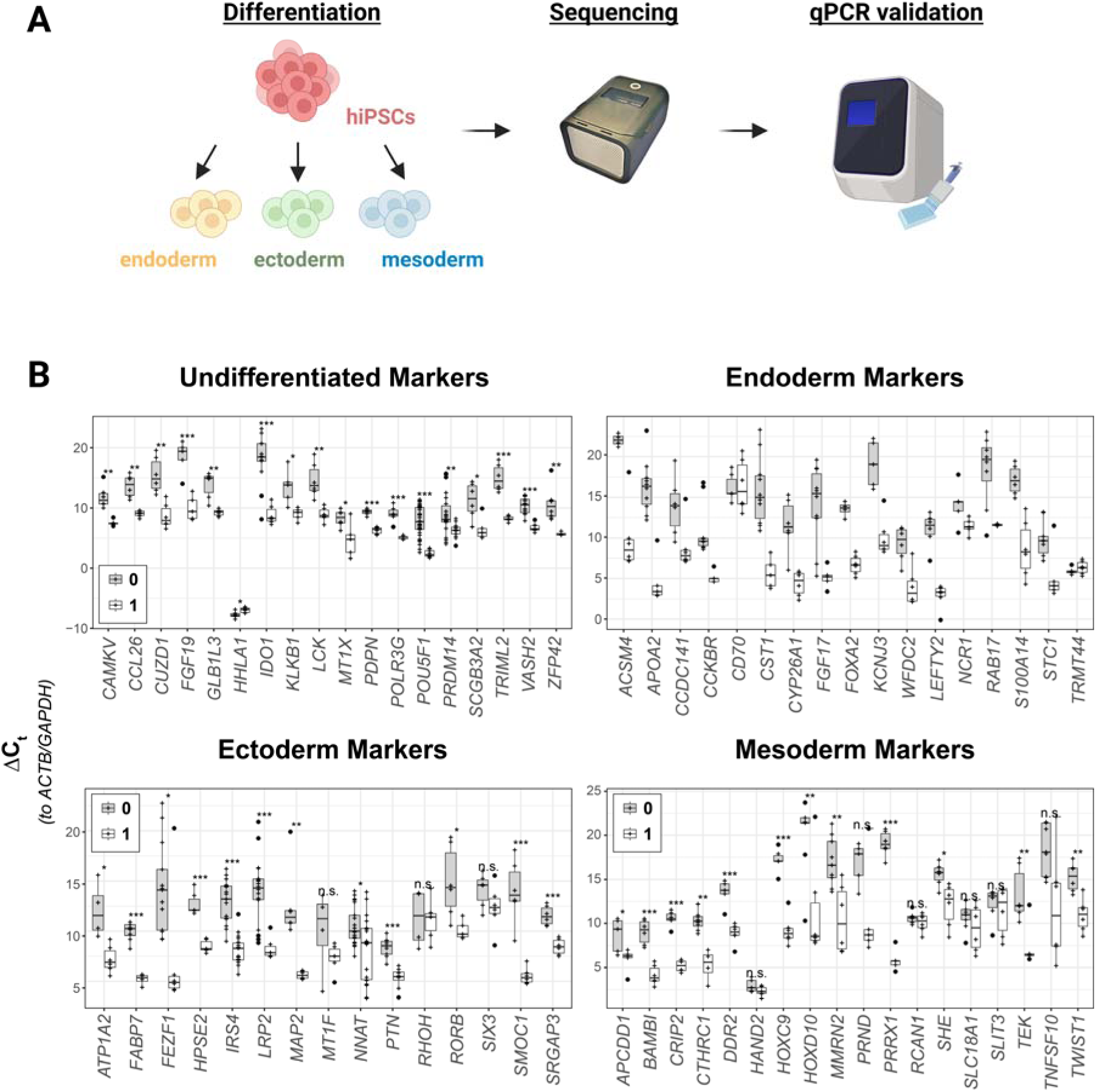
Cross-platform validation of marker genes for early hiPSC differentiation states. **(A)** Schematic overview of the validation strategy. Human induced pluripotent stem cells (hiPSCs) were maintained in the undifferentiated state or differentiated into endoderm, ectoderm, and mesoderm lineages. Candidate marker genes identified by RNA sequencing (Dobner et al., 2024a) were subsequently evaluated by qPCR in multiple independent hiPSC lines to assess their ability to discriminate between developmental states. **(B)** qPCR-based validation of candidate marker genes across early hiPSC differentiation states. A total of 68 genes were tested, including 18 undifferentiated, 17 endoderm, 15 ectoderm, and 18 mesoderm markers. Relative expression levels (normalized to *GAPDH* and *ACTB*) are shown for n ≥ 5 biological replicates. Marker performance was assessed based on enrichment in the corresponding target state compared to non-target states. Statistical significance was determined using Welch Two Sample t-tests (*p ≤ 0.05, **p ≤ 0.01, ***p ≤ 0.001; n.s., not significant).

Overall, 81% (55/68) of candidate genes were validated as robust markers capable of reliably discriminating between differentiation states, defining a set of 55 markers for early developmental stages.

To further increase robustness and expand coverage across differentiation states, we complemented this set with 11 additional markers from the original hiPSCore study RNA-seq screen (Dobner et al., 2024a) that were consistently detected in the present datasets. This resulted in a combined panel of 66 early hiPSC fate markers used for subsequent analyses (Table 1).

**Table 1.**
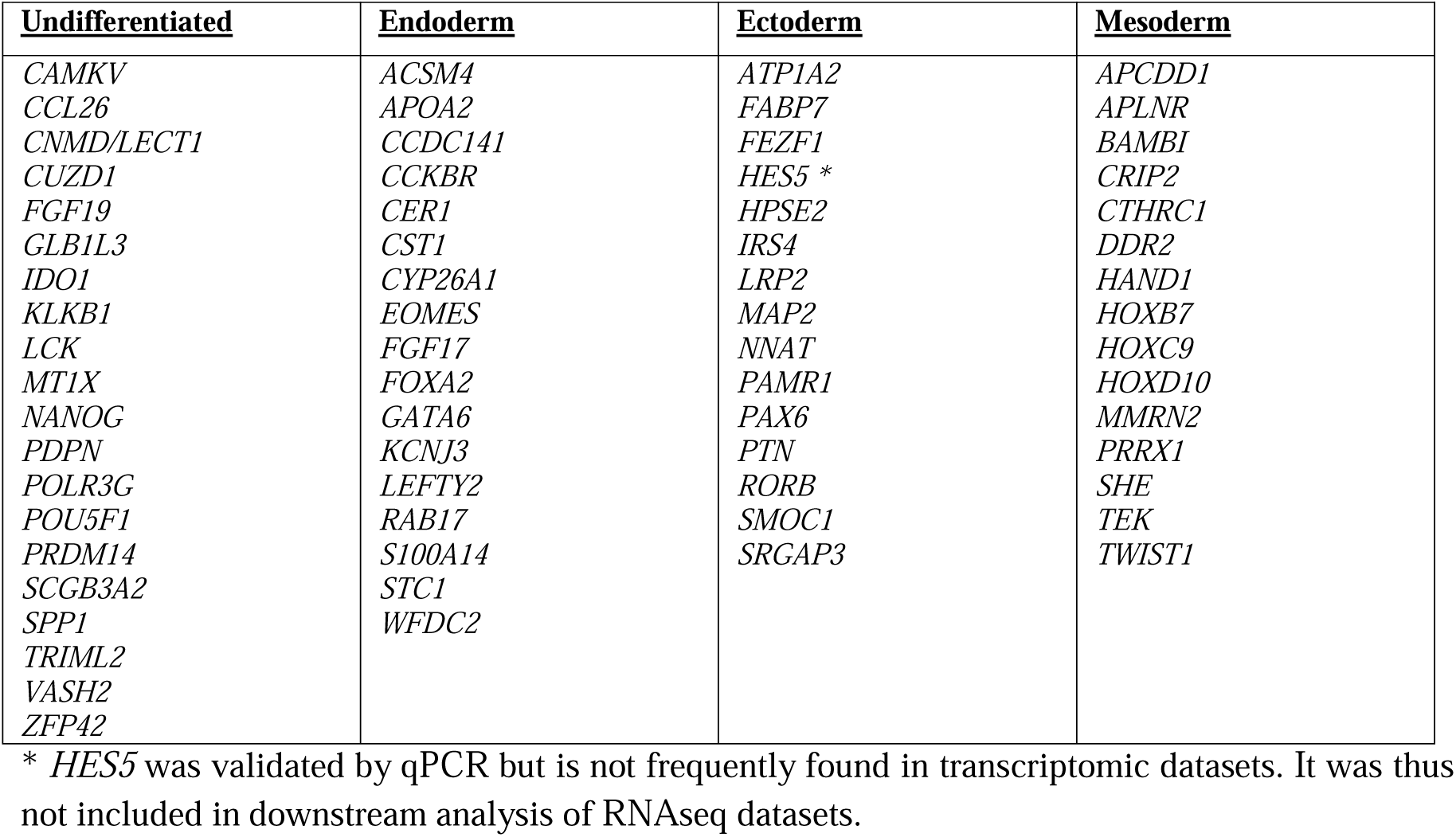
Validated marker genes to monitor undifferentiated human pluripotent stem cells and their multi-lineage differentiation obtained by directed differentiation.

Next, we asked whether these markers generalize across independent datasets and experimental contexts. To test how the validated marker set performs beyond controlled differentiation of human pluripotent stem cells, we analyzed publicly available RNA-seq datasets. We asked whether the curated markers (i) distinguish major developmental states across independent datasets, (ii) classify individual samples, (iii) capture heterogeneity within lineages, and (iv) extend to in vivo developmental contexts.

### Application of validated marker genes to assess early cell fates in public RNAseq datasets

To assess broader applicability of the validated marker gene set, we next queried publicly available RNAseq datasets deposited in the Gene Expression Omnibus (for details see Materials and Methods). We focused on datasets with available count matrices containing processed data derived from raw sequencing reads. Due to the high variety of normalization methods, we only downloaded datasets containing either raw or counts per million (cpm)-normalized counts, totaling 524 individual samples (including n = 8 from our original study). We selected undifferentiated and differentiated hiPSC- and hESC-derived samples according to the accompanying metadata and filtered for gene expression data of the core set of validated targets described above, totaling 67 early hiPSC fate markers. We performed hierarchical clustering of samples aggregated according to their annotated differentiation state by mean cpm. Intriguingly, this revealed clearly distinct patterns between differentiated samples with early germ layer-differentiated samples clustering together (**Fig. 2**). Clustering was further in line with the observed gene expression signatures of our previously published long read RNAseq samples included as separate specimen in the clustering process for validation.

**Figure 2:**
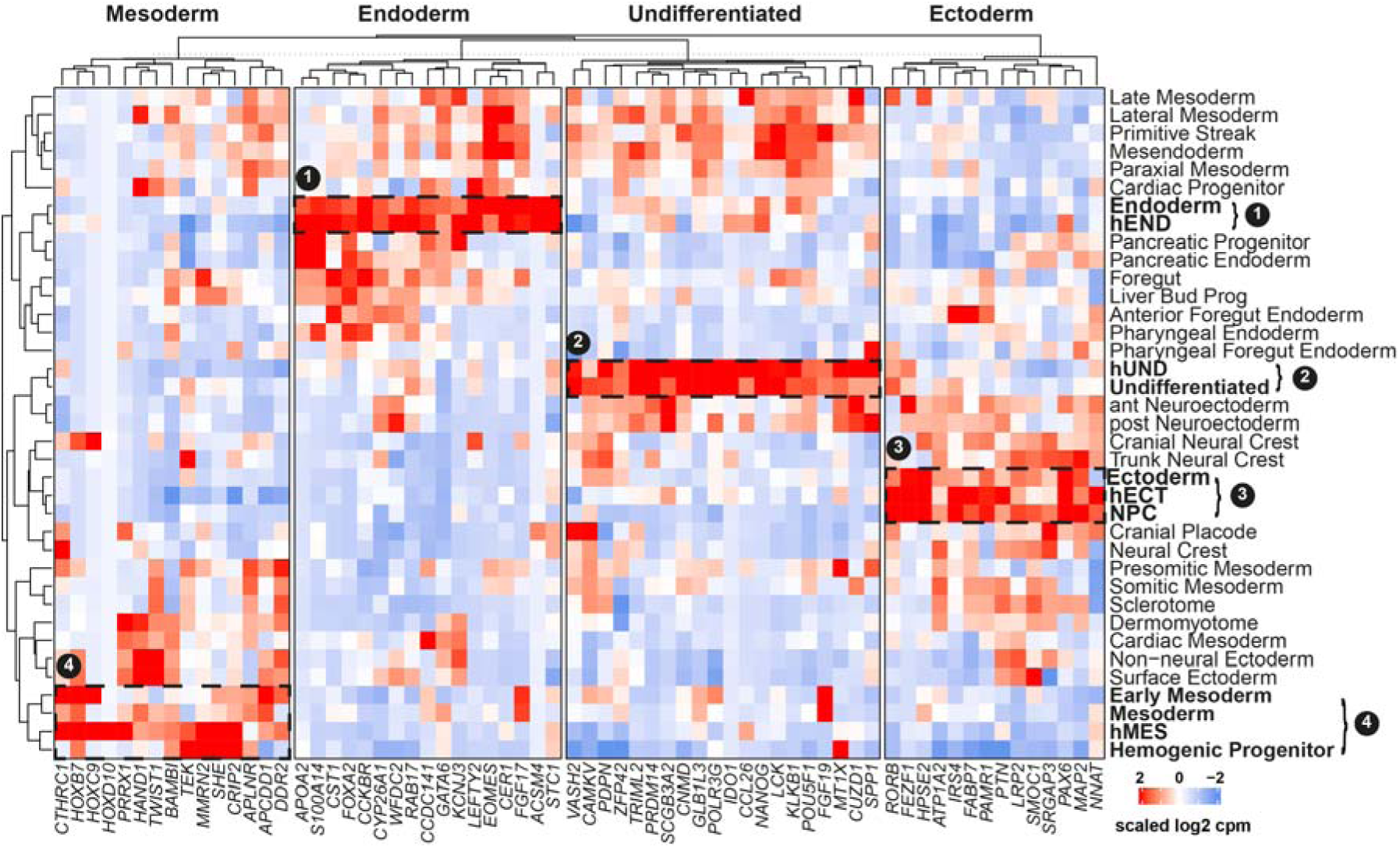
Hierarchical clustering of early human cell fates by validated marker genes. The heatmap shows early human pluripotent stem cell-related differentiation states aggregated by log2 mean counts per million (log2 cpm). Marker genes are split by allocation to one of the four early hPSC differentiation states. Rows depicting differentiation state are clustered using Pearson correlation, grouping related differentiation states in proximity, indicating sample relations. Sample sizes are indicated right next to the heatmap. hUND (hiPSCore undifferentiated), hEND (hiPSCore endoderm), hECT (hiPSCore ectoderm), and hMES (hiPSCore mesoderm) refer to the long-read RNAseq data which was originally used to identify the potential marker genes. Depicted clusters 1 – 4 indicate close sample relations: 1: hEND and aggregated Endoderm samples cluster together. 2: hUND and aggregated undifferentiated samples cluster together. 3: hECT, aggregated Ectoderm, and aggregated NPC (neural progenitor cells) samples cluster together. 4: hMES, aggregated Early Mesoderm, aggregated Mesoderm, and aggregated Hemogenic Progenitor samples cluster together. Clustering distance rows: Pearson.

These results indicate that individual differentiation states may be distinguished by the extended set of 67 core marker genes.

### Classification of individual samples

To investigate how the validated markers perform without counts averaging over all samples of a given differentiation state, we next analyzed gene expression signatures of individual samples (n = 524, Supplementary Table 3). To reduce complexity, differentiation states were grouped according to their general origin (Undifferentiated, Endoderm, Ectoderm, Mesoderm, Neural Crest, and PS + Mesendoderm). Within the Mesoderm aggregate Hemogenic Progenitor and Presomitic Mesoderm were separately labelled as they each consisted of a substantial number of samples (n = 18, n = 42, respectively). Neural progenitor cells (NPCs) were assigned to the broader (neuro)ectoderm group, whereas all samples belonging to marginal groups (n ≤ 5) were labelled as “Other”. In line with the results described before, genes of the same target cell state clustered together in hierarchical clustering (**Fig. 3**), suggesting that the validated target genes are capable of discriminating cell fates on an individual sample level. Undifferentiated state markers reliably identified and distinguished undifferentiated PSCs from other differentiation states present in the analyzed datasets. The ectoderm marker set clearly marked ectodermal cell fates and neural crest cells, whereas the endoderm marker set clearly marked endodermal cell fates. For the mesoderm marker set, a clear separation from non-mesodermal cell fates was observed, but the analyzed mesoderm data sets consisted of a more heterogenous sample group compared to the other cell fates. Primitive Streak and Mesendoderm-annotated datasets showed a mixture of undifferentiated, endoderm, and mesoderm markers, which is expected within the early differentiation dynamics of these cells.

**Figure 3:**
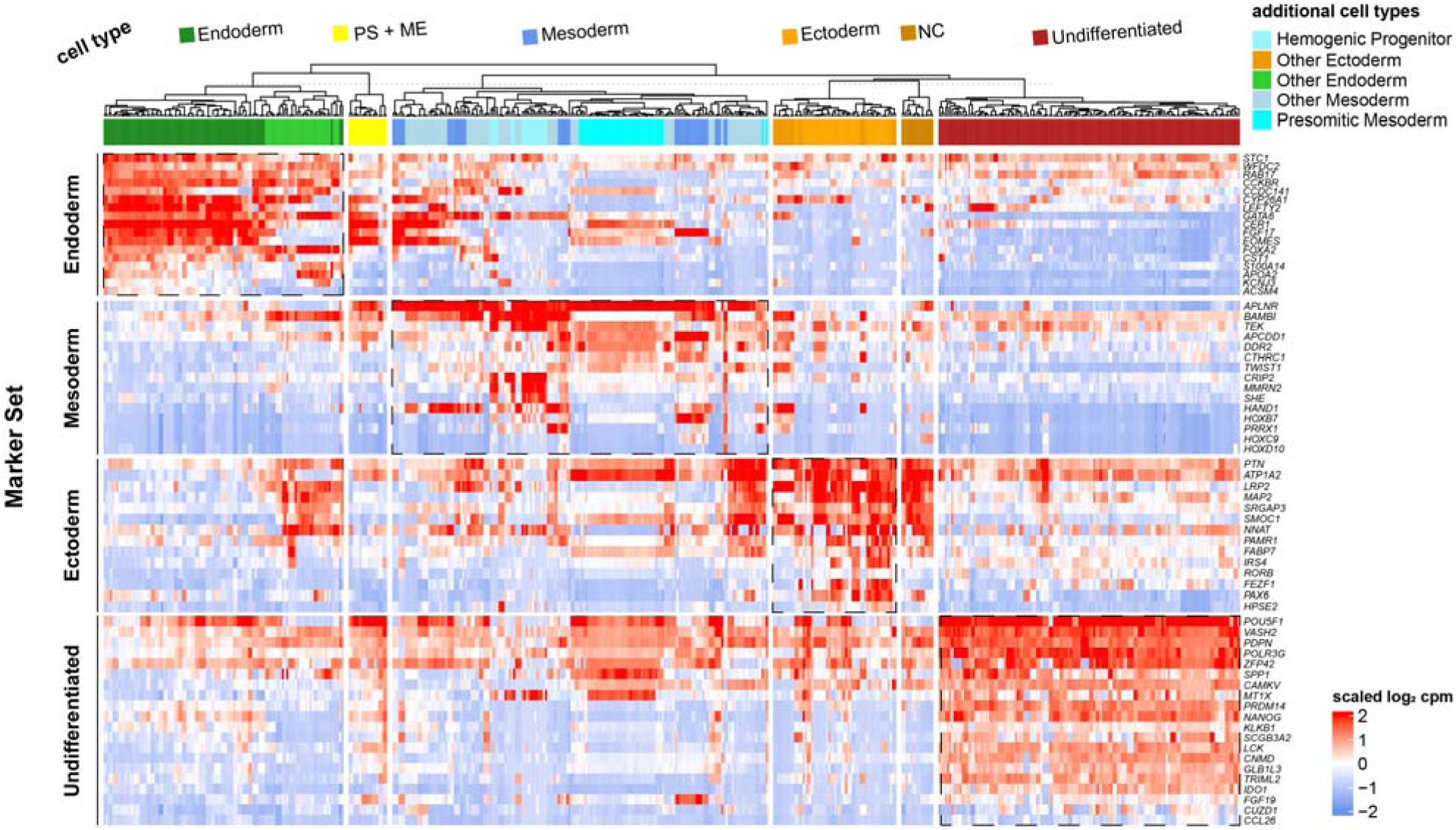
Hierarchical clustering of early human pluripotent stem cell-derived germ layer differentiation states. The heatmap shows gene expression values of indicated genes as scaled log2 counts per million (log2 cpm). The heatmap is horizontally split by genes allocated to a specific target differentiation state (undifferentiated and germ layer-differentiated human pluripotent stem cells), and vertically by cell type. For clarity, individual cell types were aggregated into the broader categories of “Mesoderm”, “Undifferentiated”, “Ectoderm”, “Endoderm”, “Neural Crest”, and “Primitive Streak (PS) + Mesendoderm”. Additional cell types are indicated in the respective broader group by color-coding. Dashed borders indicate matching cell types and marker gene sets. Clustering indicates that the validated marker genes are able to distinguish different cell types based on expression signatures in the analyzed datasets. PS: primitive streak, ME: mesendoderm. Clustering distance: Pearson.

When restricting the clustering more strictly to the early germ layer differentiation fates, the actual target cell fates of the original publication, the pattern became even clearer (**Fig. S1, Fig. 4**), with high within early cell fate correlations (Endoderm r□=0.810; Undifferentiated r□=0.796; Ectoderm r□=0.706; Mesoderm r□=0.528). Between early cell fates, Endoderm and Ectoderm (r□=−0.029), and Undifferentiated-Mesoderm (r□=0.037) were uncorrelated, while Endoderm-Mesoderm (r□=0.167) were mildly, and Ectoderm-Undifferentiated were moderately (r□=0.384) correlated **(Fig. 4)**. This pronounced heterogeneity within mesodermal samples, raised the question of whether this reflects biological substructures or loss of lineage identity in annotated sample data. As within sample correlation among the analyzed cell fates were lowest, we analyzed a subset of mesoderm-differentiated samples of four independent hiPSC lines by flow cytometry. Notably, we found differing percentages of CD144^+^ endothelial-biased, and CD144^-^non-endothelial mesoderm **(Fig. S2)**, indicating that mesoderm differentiation in general is more heterogenous than endoderm and ectoderm differentiation.

**Figure 4:**
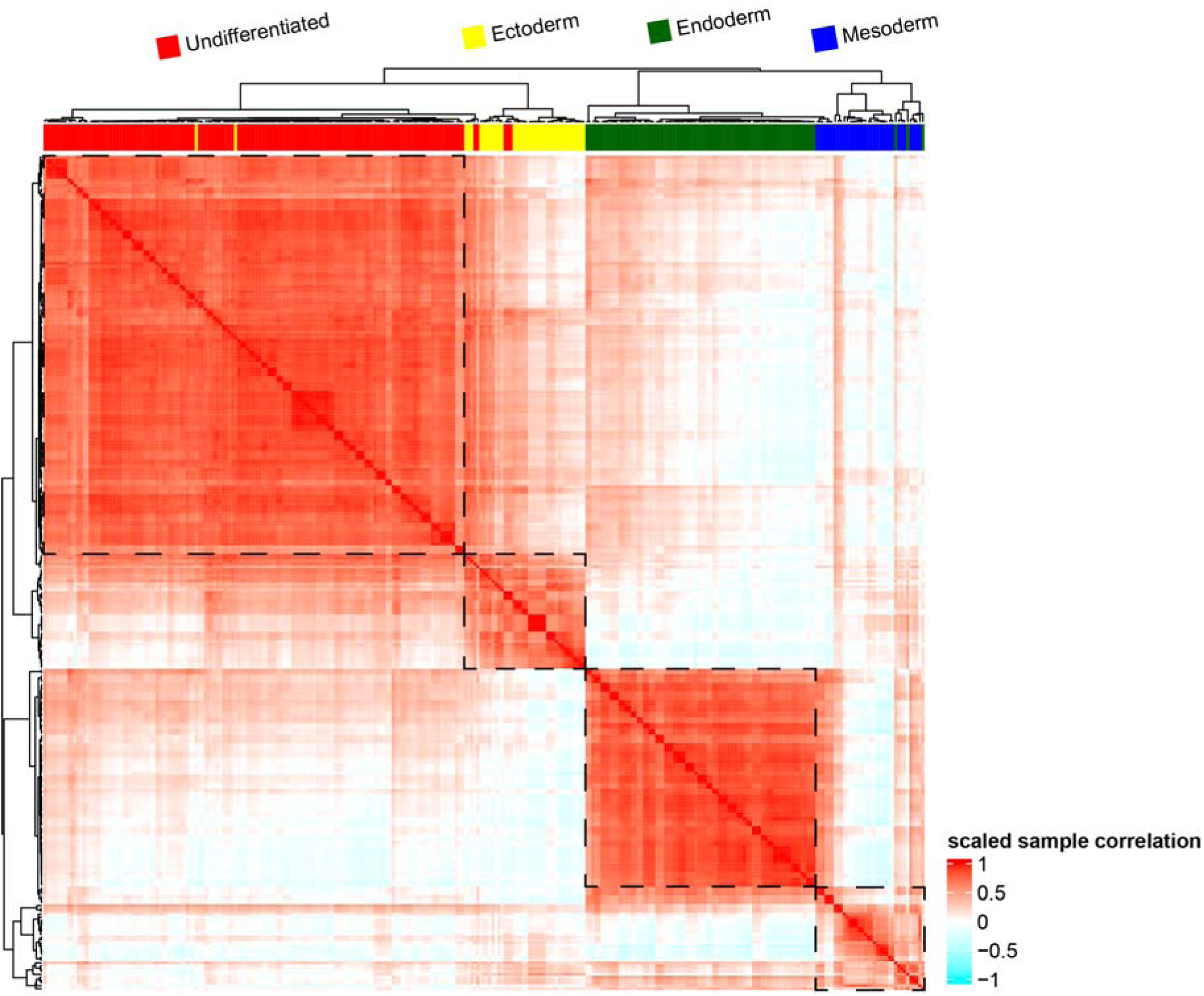
Hierarchical clustered correlation heatmap of early human pluripotent stem cell-derived germ layer differentiation states. The heatmap is based on calculated sample correlations between individual samples of each of the four early human induced pluripotent stem cells (hiPSCs) differentiation states. Each tile represents one of 523 pairwise correlations. Sample correlations have been scaled and are represented hierarchically clustered to indicate high sample correlations. Clear patterns of undifferentiated, ectoderm, endoderm, and mesoderm individual samples are apparent. Only few (n < 5) individual samples cluster in disagreement with their cell fate annotation. Black dashed rectangles indicate distinctly separatable cell fate clusters. Clustering distance: Pearson.

### Subclassification of early developmental lineages

We further explored whether the validated marker set enables finer subclassification of early germ layer derivatives (**Fig. S3**). Endoderm samples showed overlap with mesendodermal samples, but were clearly separable from later lineages using the extended validated endoderm marker gene set (**Fig. S3A)**, while samples aggregated to ectoderm (NPC, neuroectoderm) showed two distinct neuroectodermal clusters largely separate from other cell types (**Fig. S3B**). Mesoderm samples split into more distinct stages (early, late, somitic, presomitic, lateral) alongside other mesoderm-related samples revealed distinct clustering for presomitic mesoderm, mesoderm, sclerotome, and early mesoderm, while other mesoderm-related samples were more similar based on clustering (**Fig. S3C**).

### Validated marker set labels cell types in single cell RNAseq datasets of human embryo

Developing organisms comprise dynamic cell states rather than static, pure populations. To evaluate our marker framework in this context, we applied it to a scRNA-seq dataset of a gastrulating human embryo (Zhao et al., 2025) containing annotated data from Carnegie Stage 7 (day 16–19; Tyser et al., 2021)). Due to the sparsity of scRNA-seq data relative to bulk RNA-seq, we aggregated our markers into four module scores representing the undifferentiated state (“Undiff”) and the three germ layers (“Endo”, “Ecto”, “Meso”). We visualized the dataset’s thirteen annotated cell types (**Fig. 5A**) and projected the module scores onto the UMAP embedding (**Fig. 5B**).

**Figure 5:**
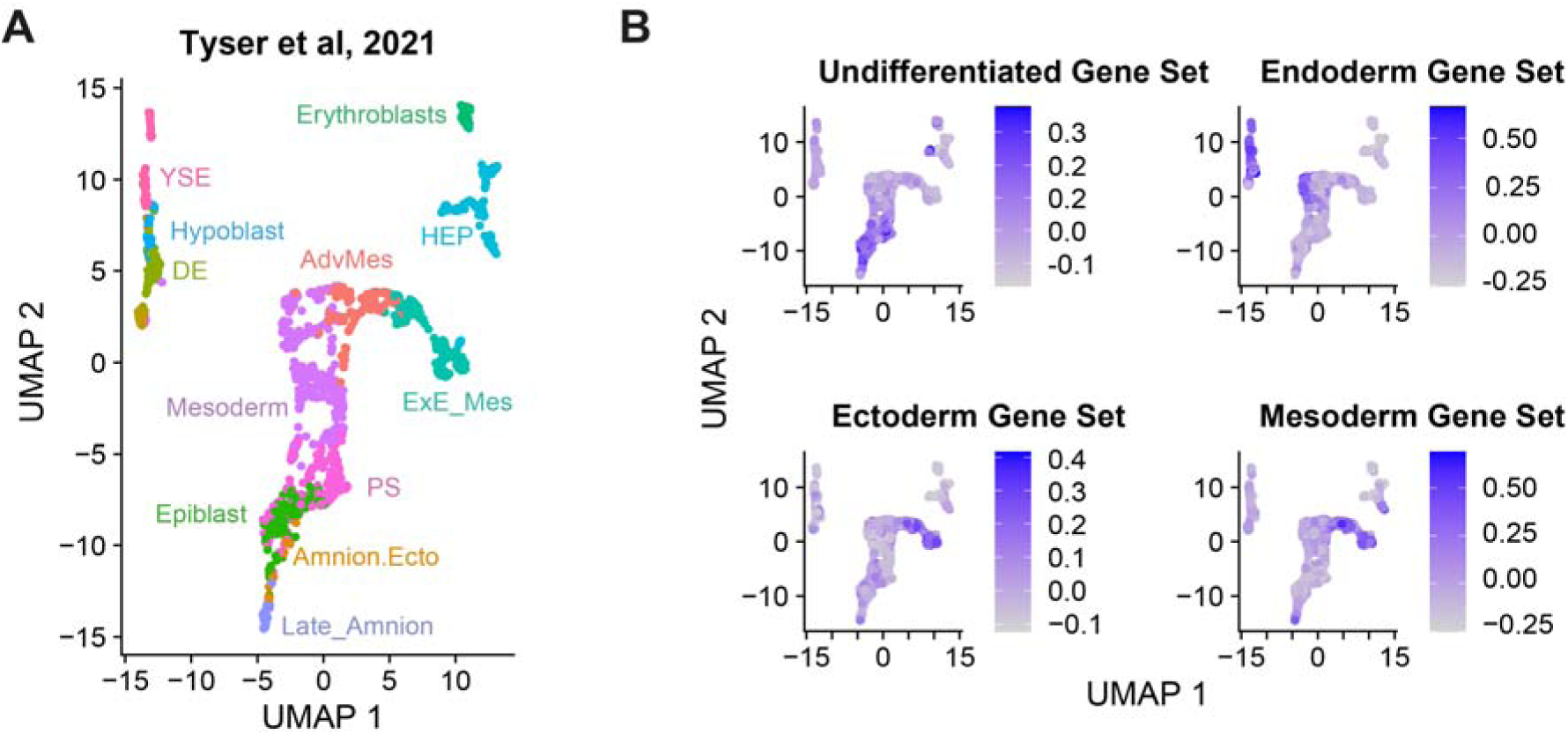
Analysis of validated marker genes in single cell RNA sequencing reference dataset of a gastrulating human embryo. (A) The reference scRNAseq dataset was derived from Tyser et al, 2021 and contains 13 distinctly annotated cell types of a Carnegie Stage 7 gastrulating human embryo. (B) Validated marker genes were aggregated into “Module Scores” according to their target cell fate allocation (Undifferentiated, Endoderm, Ectoderm, and Mesoderm Gene Set) and relative gene expression scores mapped to the annotated cell types of the reference dataset. Annotations were derived from the processed dataset. AdvMes: Advanced Mesoderm, Amnion.Ecto: Amniotic Ectoderm, Axial Mes: Axial Mesoderm, DE: Definitive Endoderm, ExE_Mes: Extryembryonic Mesoderm, HEP: Hemato-Endothelial Progenitors, PriS: Primitive Streak, YSE: Yolk Sac Endoderm.

The “Undiff” score was enriched primarily in the epiblast (**Fig. S4A**), validating its utility for identifying pluripotent cells. The “Endo” score identified yolk sac endoderm, hypoblast, definitive endoderm, and mesoderm (Fig. S5 B), demonstrating robustness in dynamic conditions. The “Ecto” score mapped to extraembryonic mesoderm (Fig. S5 C); this aligns with the neural groove developing around CS8 (O’Rahilly and Gardner, 1979; O’Rahilly and Müller, 1981), consistent with the longer differentiation trajectory required for neuroectoderm. Finally, the “Meso” score successfully labeled mesodermal populations, including late amnion and advanced mesoderm (Fig S5 D), confirming the gene set’s specificity in this complex tissue.

### Linking gene expression levels to protein abundance

Because developmental identity is ultimately executed at the protein level, we examined whether our transcript-based marker signatures are reflected in protein abundance. While mRNA and protein levels are generally correlated, differential gene expression does not always translate to altered protein levels (Buccitelli and Selbach, 2020). Although RNA-seq remains the standard for analyzing global cellular dynamics due to its accessibility and ease of interpretation, validating transcriptomic findings at the protein level is crucial for understanding functional cellular responses. However, unbiased proteomics datasets remain less common in hiPSC research than RNA-seq datasets (Lindoso et al., 2019).

To address this, we performed ultra-deep proteomic (> 7,000 proteins per sample on average) profiling by mass spectrometry (MS) on the hiPSC reference line KOLF2.1J, analyzing triplicates of undifferentiated cells and cells differentiated into the three germ layers (**Fig. 6A**). This approach extended our recent metabolomic characterization of this line (Dobner et al., 2024b). Differentially abundant proteins were identified using empirical Bayes statistics (see Materials and Methods).

**Figure 6:**
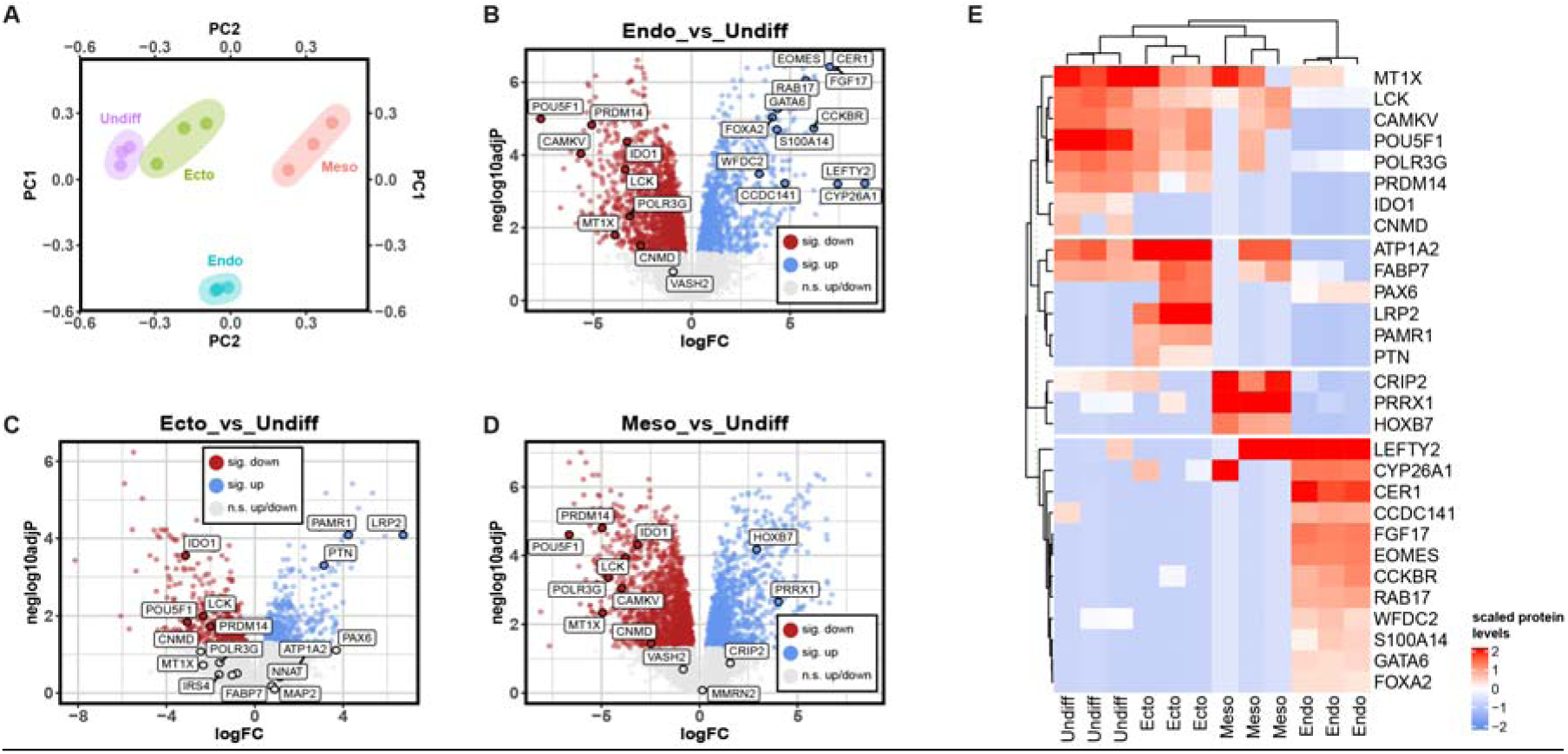
Proteome profiling of undifferentiated and trilineage-differentiated human induced pluripotent stem cells by mass spectrometry. (A) Biological triplicates of undifferentiated, endoderm-, ectoderm-, and mesoderm-differentiated KOLF2.1J human induced pluripotent stem cells (hiPSCs) were subjected to mass spectrometry. Principal Component (PC) analysis after data processing and missing value imputation demonstrates clear separation between distinct differentiation states. (B) Volcano plot showing differentially abundant proteins between endoderm and undifferentiated hiPSCs. RNAseq- and qPCR-validated undifferentiated state and endoderm markers are labelled. (C) Volcano plot showing differentially abundant proteins between ectoderm and undifferentiated hiPSCs. RNAseq- and qPCR-validated undifferentiated state and ectoderm markers are labelled. (D) Volcano plot showing differentially abundant proteins between mesoderm and undifferentiated hiPSCs. RNAseq- and qPCR-validated undifferentiated state and mesoderm markers are labelled. (B – D) Proteins are labelled as points in red (significantly less abundant in differentiated vs undifferentiated state), blue (significantly more abundant in differentiated vs undifferentiated state), and grey (no significantly different abundance). Statistical significance was inferred at an “Benjamini Hochberg” post-hoc adjusted p value of ≤ 0.05. (E) Clustered heatmap visualizing genes identified by long read sequencing as protein levels of mass spectrometry (MS) data in early germ layer differentiated and undifferentiated human induced pluripotent stem cells (hiPSCs) of the KOLF2.1J line. n.s.: not significant. logFC: log fold change.

Due to the inherent sensitivity limits of mass spectrometry, not all transcript-defined markers were detectable at the protein level, and subsequent analyses were therefore restricted to the subset of markers with measurable protein abundance.

We assessed the concordance between our validated marker panel and protein abundance by visualizing comparisons between undifferentiated cells and germ layers, as well as cross-lineage comparisons, using volcano plots (**Figs. 6B–C, S5A–C**).

Most validated gene markers showed consistent changes in protein abundance. Upon differentiation, 100% (12/12) of detected endoderm proteins (Fig. 6B) and 75% (3/4) of mesoderm proteins (**Fig. 6D**) were significantly enriched (adjusted p ≤ 0.05). For ectoderm, 33% (3/9) of proteins were significantly enriched (**Fig. 6C**), with PAX6 approaching significance (p = 0.078). Conversely, undifferentiated markers were significantly downregulated in 89% of cases compared to endoderm and mesoderm, but only 57% compared to ectoderm (**Figs. 6B–D**). This latter finding reflects the closer transcriptional proximity of undifferentiated cells and ectoderm observed in the PCA analysis (**Fig. 6A**).

In cross-lineage comparisons, endoderm markers were consistently enriched, showing significant increases in 100% (12/12) of cases relative to both ectoderm and mesoderm (**Figs. S5A–B**). Ectoderm markers were significantly enriched compared to mesoderm in 83% (5/6) of cases (Fig. S6C), but significantly decreased compared to endoderm in 56% (5/9) of cases (**Fig. S5A**). Similarly, mesoderm markers were significantly decreased compared to endoderm (75%, 3/4) and ectoderm (100%, 3/3) (**Figs. S5A, S5C**). Taken together, these results demonstrate high congruence between markers identified by RNA-seq, cross-validated by qPCR, and protein levels detected by MS (**Fig. 6E).**

### DeepDiff: An interactive resource for gene and protein expression analysis

To facilitate the practical application of our validated marker framework, we translated our findings into an accessible analysis resource. We therefore developed “DeepDiff”, an R Shiny-based web application (http://jdobner.shinyapps.io/diff_sets) that allows for the rapid assessment of gene expression across differentiation states. We aggregated CPM-normalized reads to calculate descriptive statistics (mean, median, range, standard deviation, and quartiles) for 33 distinct cell fates as annotated by the original dataset authors. DeepDiff provides visual and tabular representations of these data (**Fig. 7A**), enabling users to identify candidate marker genes and assess their functional relevance for specific lineages. Beyond data visualization, we implemented a supervised machine learning model for classifying undifferentiated cells and early germ layers using CPM-normalized RNA-seq data (**Fig. 7B**). We also integrated the MS proteomics data via a dedicated “Protein” tab, allowing for direct comparison of protein abundance across the four early differentiation states (**Fig. 7C**). Finally, to bridge transcriptomic and proteomic layers, we included a correlation analysis module based on scaled gene expression and MS intensity measurements (**Fig. 7D**). These features provide a comprehensive platform for exploring the relationship between gene expression and protein levels, aiding in the planning and execution of differentiation experiments.

**Figure 7:**
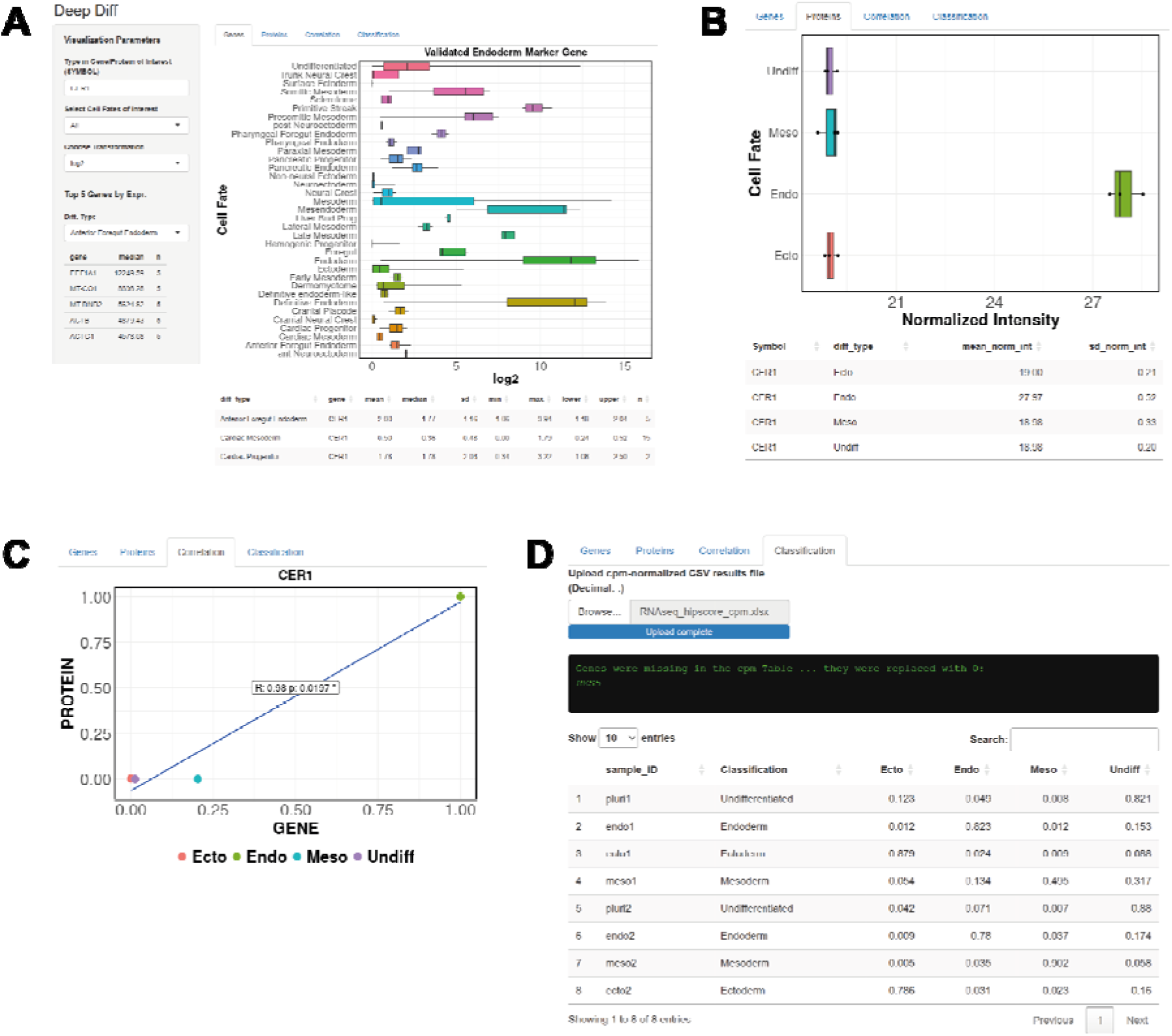
Rapid assessment and linkage of gene expression and protein levels using DeepDiff. A: DeepDiff is a R Shiny-based web resource enabling users to access gene expression values of RNAseq datasets containing early pluripotent stem cell-derived differentiation states without the need for bioinformatic expertise. The Gene Symbol of interest is typed into a text field and gene expression values are displayed as counts per million (cpm), log2 cpm, or sqrt-transformed cpm. Either all (All) annotated cell fates or a subset (Early Germ Layers, Endodermal, Ectodermal, Mesodermal) can be visualized simultaneously. The top 5 (by cpm value) expressed genes are displayed and aid to evaluate, e.g., whether a gene of interest is present/absent in a given cell type. To reduce file size and enable web hosting, statistics were precomputed and are displayed in a table below the boxplot visualization. B: DeepDiff enables assessment of protein levels determined by MS in early germ layer differentiated and undifferentiated hiPSCs. The CER1 protein is exemplarily shown and found in endoderm-differentiated hiPSCs at the highest level. C: The correlation between gene expression levels and protein levels is an integral part of the DeepDiff web resource. It enables quick assessment of how gene expression levels correlate with protein levels. Scaled gene expression (counts per million) and normalized MS-derived intensities of aggregated of samples/proteins are displayed and a simple correlation statistic calculated which can give insights into gene expression dynamics. VEGFA is shown as an example. D: DeepDiff offers a machine learning-based classification of early human pluripotent stem cell germ layer samples based on RNAseq-derived cpm-normalized gene expression values. The hiPSCore (Dobner et al, 2024) dataset is exemplarily shown.

## Discussion

Early human development is characterized by continuous and dynamic cell identity transitions. This complexity presents the challenge of distinguishing stable lineage commitment from the graded modulation inherent to developmental trajectories (Boroviak and Nichols, 2017; Mittnenzweig et al., 2021; Pijuan-Sala et al., 2019). Here, we establish and validate a transferable marker framework that addresses this challenge by robustly defining early developmental identity across platforms, experimental systems, and molecular layers.

Previous marker-based approaches have primarily assigned discrete fate labels within narrowly defined experimental contexts (Kiselev et al., 2019; Morris, 2019; Trapnell, 2015). While effective for identifying major lineage transitions, these approaches often fail to resolve early developmental states that are heterogeneous or shaped by perturbation. Our results demonstrate that early developmental identity can be defined using markers that are both robust to technical variability and sensitive to biological modulation. Notably, we show that lineage identity remains detectable despite substantial transcriptional heterogeneity, particularly within the mesoderm lineage, highlighting the framework’s ability to capture stable identity without collapsing nuanced intra-lineage variation.

To date, marker validation in the hPSC field has focused predominantly on the undifferentiated state (Andrews and Gokhale, 2024; Ramirez et al., 2011; Sekine et al., 2020), with germ-layer markers receiving less systematic attention. The ISSCR and related initiatives provide marker recommendations for trilineage assessment (Allison et al., 2018; Bock et al., 2011), but these recommendations are rooted in historical convention rather than systematic, evidence-based curation across platforms. Our data confirm that a subset of commonly used markers lacks the specificity to unambiguously resolve early trilineage differentiation states (Dobner et al., 2024a; Kuang et al., 2019), underscoring the need for rigorous, cross-platform validation of markers recommended by standard-setting bodies. Building on our previous hiPSCore (Dobner et al., 2024a), we consolidated a curated panel of 67 marker genes. The generalizability of this panel across diverse sequencing devices, protocols, and laboratories, together with its ability to label cell types in in vivo human embryo datasets and concordant protein-level expression for a subset of detectable markers, supports its broad utility.

We therefore propose that these validated markers be considered for integration into standardized guidelines for screening early differentiation states, addressing a gap in current recommendations that lack evidence-based, cross-platform validated marker panels for directed trilineage differentiation.

Notably, discrepancies between RNA-seq and qPCR measurements likely reflect both technical and biological factors, including assay-specific sensitivities and transcript-level variability, underscoring that transcript abundance does not translate in a one-to-one manner across platforms and highlighting the importance of systematic experimental validation. Our framework bridges this gap by rigorously filtering candidates through both sequencing and targeted validation, ensuring that the final marker set is practically applicable for routine quality control.

To maximize the utility of these findings, we translated our framework into DeepDiff, a publicly accessible web resource. DeepDiff serves as a bridge between complex multi-omic datasets and the end-user, requiring no bioinformatic expertise to operate. It provides: (1) rapid assessment of gene expression across 33 hPSC differentiation states; (2) access to proteomic data for the reference line KOLF2.1J, filling a gap in available protein-level resources; (3) correlation analysis linking transcript and protein dynamics; and (4) a machine learning classifier for the automated annotation of bulk RNA-seq data. By integrating curated markers, multi-omic validation, and accessible tools, this work establishes a standardized foundation for studying early human developmental identity and its modulation by genetic or environmental perturbations.

### Limitations and future directions

While our curated marker panel demonstrates robust performance, the yet uncharacterized part of the initial pool of 172 candidate genes likely contains additional informative markers that could further expand and refine the framework. In addition, the mass spectrometry data highlight protein-level candidates that warrant further functional and validation studies. Future work will focus on systematically integrating these candidates to enhance lineage resolution and robustness. In parallel, DeepDiff can be further improved by incorporating additional datasets, including emerging single-cell and spatial transcriptomic data, expanding protein-level validation, and refining its classification algorithms to support more complex and heterogeneous developmental states. While additionally to the Tyser dataset, spatial transcriptomics datasets of a CS8 human embryos useful for this study supposedly exist (Xiao et al., 2024), we were unable to make use of these data due to unavailable Stereo-seq chip masks.

## Supplementary Information

**Figure S1:**
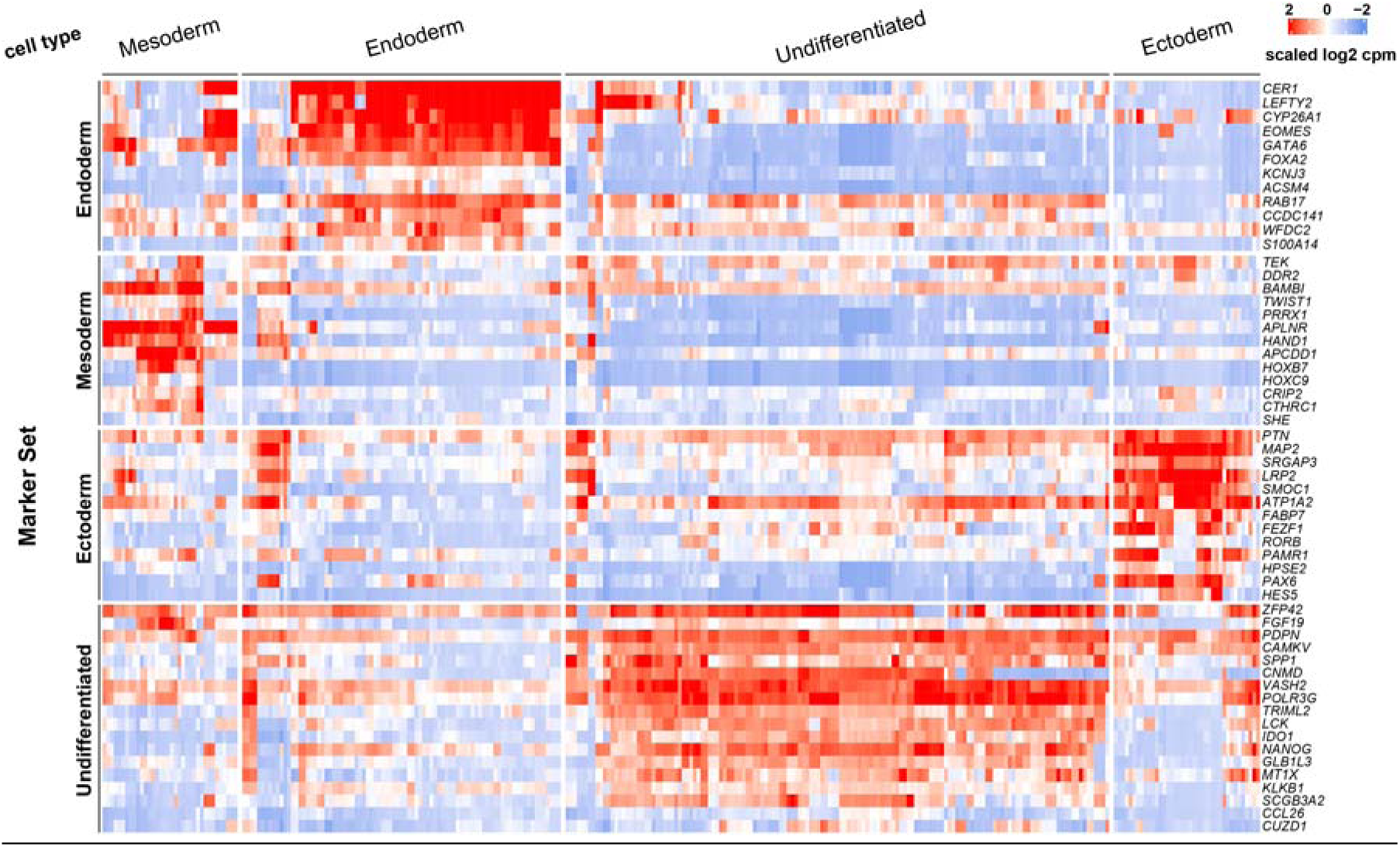
Hierarchical clustering of human pluripotent primary germ layer states and undifferentiated samples. The heatmap shows gene expression values of indicated genes as scaled log2 counts per million (log2 cpm). The heatmap is horizontally split by genes allocated to a specific target differentiation state (undifferentiated and germ layer-differentiated human pluripotent stem cells), and vertically by cell type. Clustering indicates that the validated marker genes are able to distinguish different cell types based on expression signatures in the analyzed datasets. Clustering distance: Pearson.

**Figure S2:**
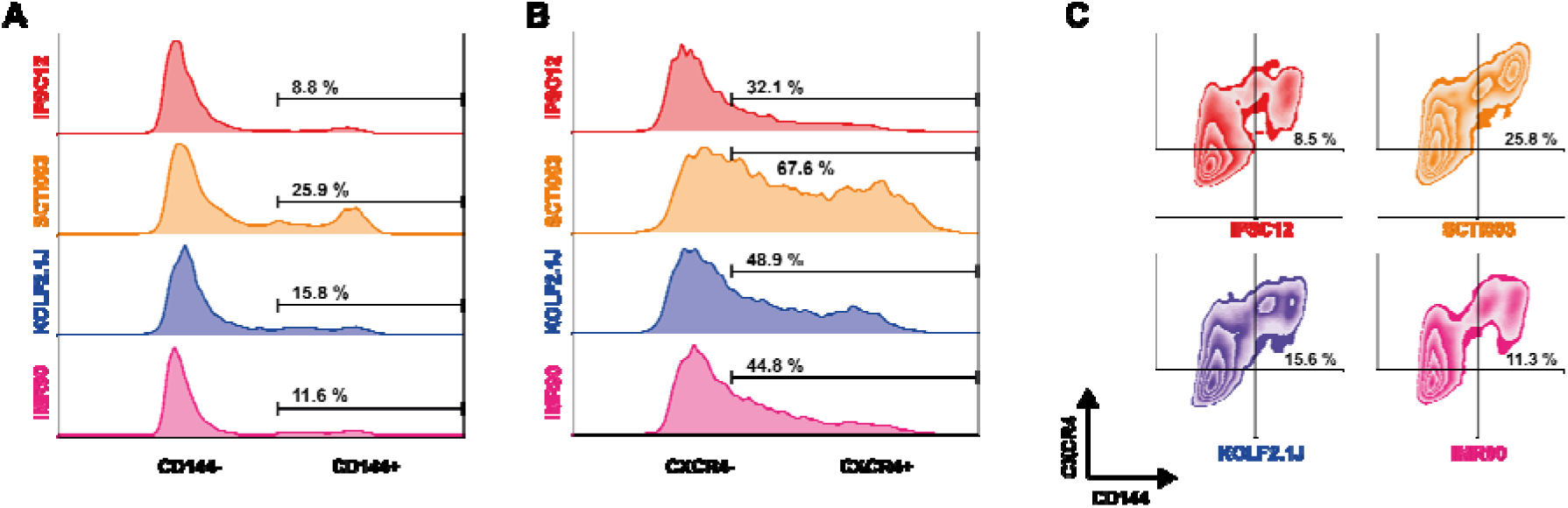
Flow cytometry analysis of mesoderm-differentiated human induced pluripotent stem cells. Four independent human induced pluripotent stem cell (hiPSC) lines were differentiated into mesoderm using a commercially available kit. (A) CD144-positive (+) endothelial mesoderm cells range from 8 to 25 %. (B) CXCR4-positive (+) mesodermal cells range from 32 to 67 %. (C) CD144/CXCR4 double-positive cells range from 8 to 25 %, indicating that CXCR4 is not specific to endothelial mesoderm.

**Figure S3:**
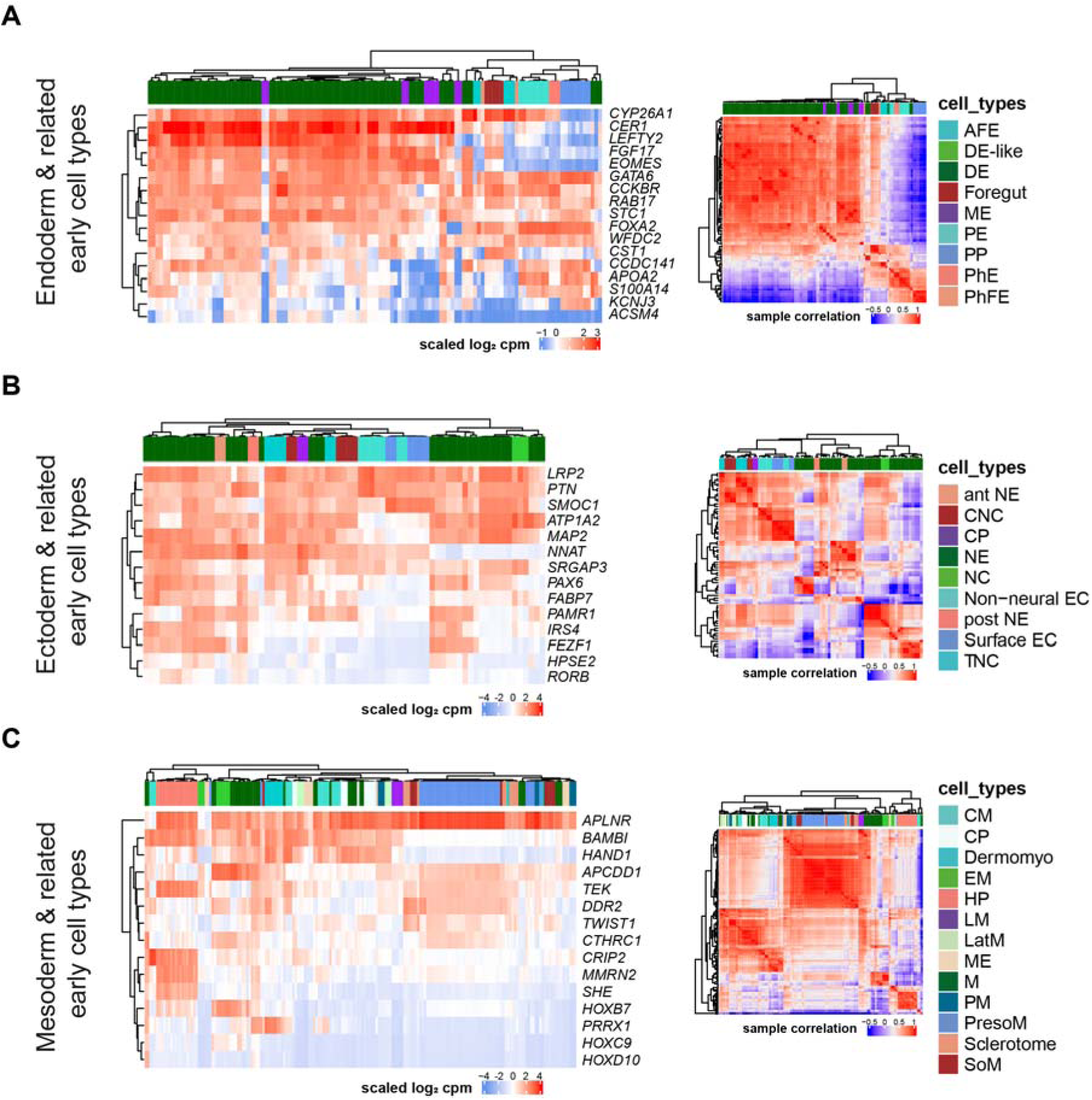
Hierarchical clustering of distinct early germ layer states and related lineages. Individual Heatmaps display sample relations of early human pluripotent stem cell germ layer fates and related downstream lineages clustered by gene scaled log2 counts per million (cpm-)normalized gene expression values of the respective germ layers (left) and visualized as pairwise correlations (right) based on the same gene expression signatures. (A) Definitive Endoderm clusters separately based on extended hiPSCore gene expression signatures compared to other endoderm-related lineages. Minimal overlap with mesendodermal samples can be assumed from the heatmaps. (B) Two distinct Neuroectoderm blocks interspersed with anterior Neuroectoderm and neural crest samples cluster separately to other ectoderm-related cell fates based on gene expression signature and correlation-based clustering. (C) Mesoderm, sclerotome, and presomitic mesoderm samples cluster separately, while other mesoderm-related cell types share overlapping gene expression signatures, indicating that Mesoderm gene expression signatures might be shared more broadly among mesodermal lineages. Clustering distance: Pearson. AFE: Anterior Foregut Endoderm; DE: Definitive Endoderm; ME: Mesendoderm; PE: Pancreatic Endoderm; PP Pancreatic Progenitor; PhE: Pharyngeal Endoderm; PhFE: Pharyngeal Foregut Endoderm; NE: Neuroectoderm; CNC: Cranial Neural Crest; CP: Cranial Placode; NC: Neural Crest; EC: Ectoderm; TNC: Trunk Neural Crest; CM: Cardiac Mesoderm; CP: Cardiac Progenitor; Dermomyo: Dermomyotome; EM: Early Mesoderm; HP: Hemogenic Progenitor; LM: Late Mesoderm; LatM: Lateral Mesoderm; ME: Mesendoderm; M: Mesoderm; PM: Paraxial Mesoderm; PresoM: Presomitic Mesoderm; SoM: Somitic Mesoderm.

**Figure S4:**
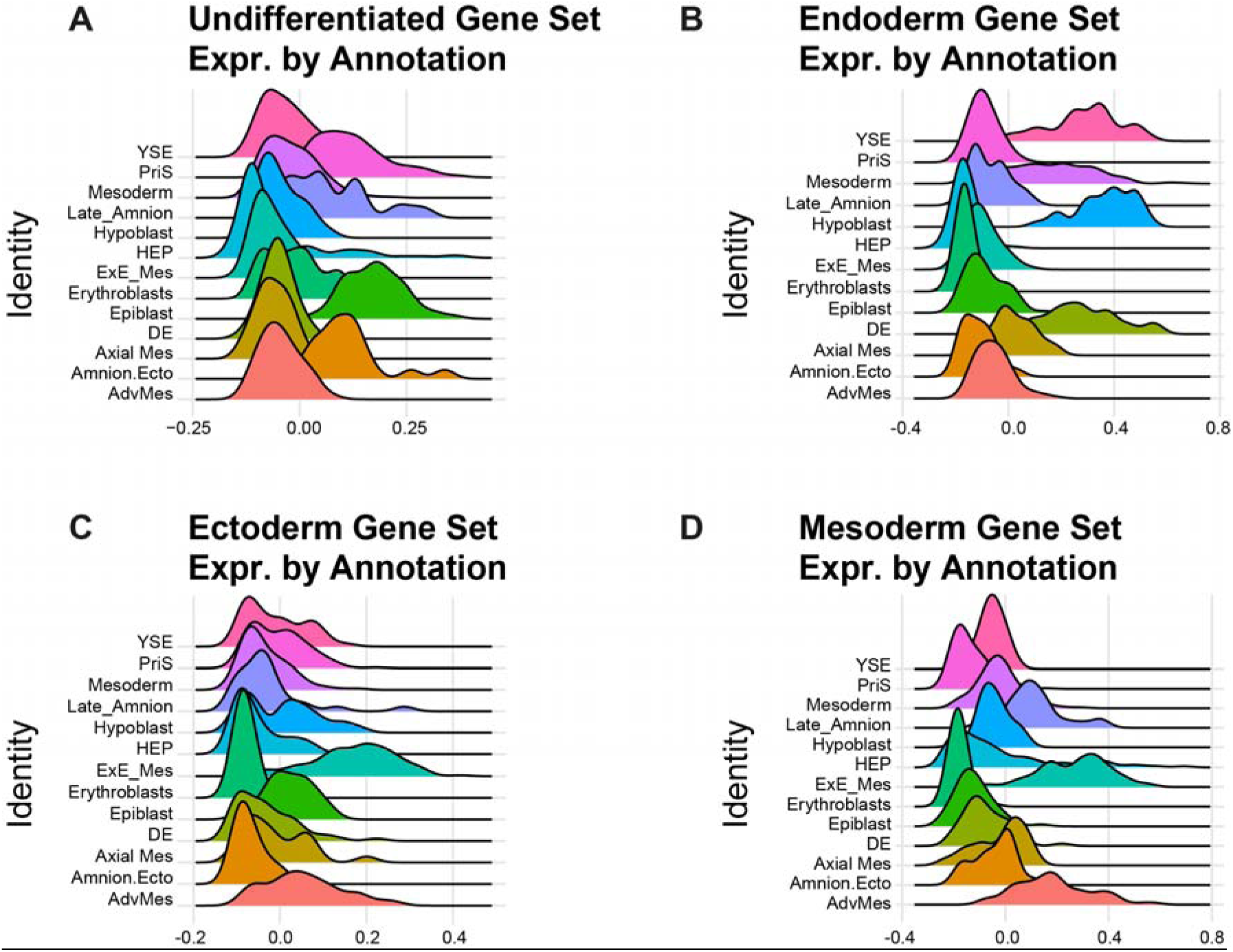
Ridge plots indicating marker gene-derived Module Score expression across human gastrulating Carnegie Stage 7 embryo. Validated marker genes were aggregated into “Module Scores” according to their target cell fate allocation (A: Undifferentiated, B: Endoderm, C: Ectoderm, and D: Mesoderm Gene Set). Relative gene expression scores mapped to the annotated cell types of the reference dataset are displayed as ridge plots to visualize enrichment of Module Scores across distinct differentiation states. Annotations were derived from the processed dataset. AdvMes: Advanced Mesoderm, Amnion.Ecto: Amniotic Ectoderm, Axial Mes: Axial Mesoderm, DE: Definitive Endoderm, ExE_Mes: Extryembryonic Mesoderm, HEP: Hemato-Endothelial Progenitors, PriS: Primitive Streak, YSE: Yolk Sac Endoderm.

**Figure S5:**
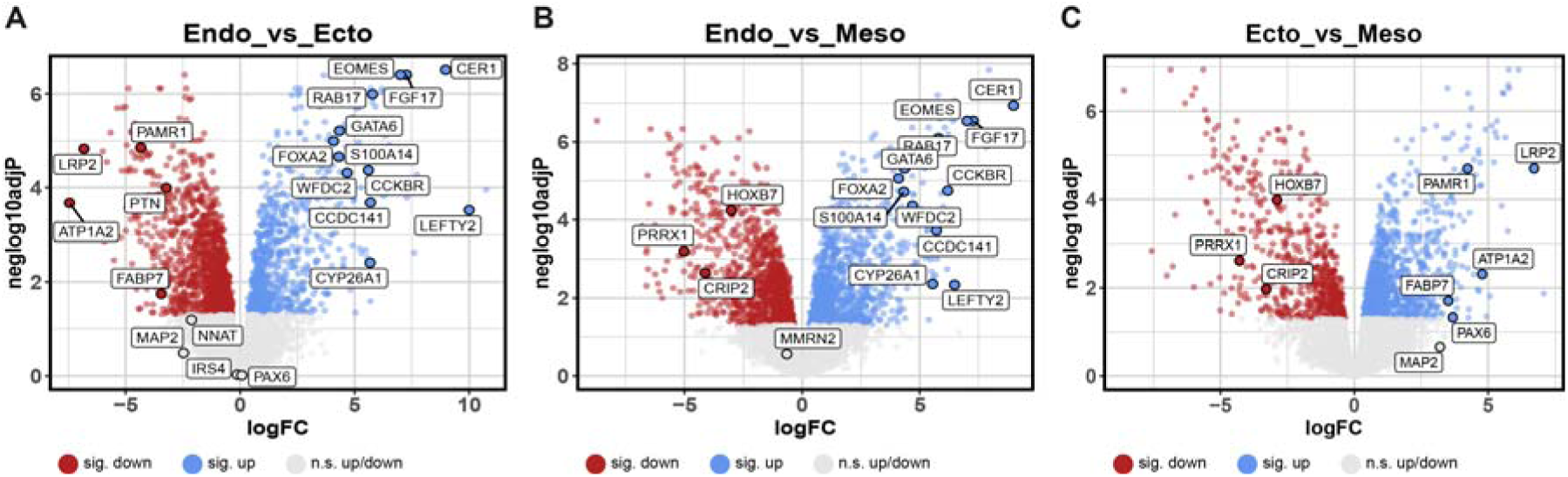
Proteome comparison of trilineage-differentiated human induced pluripotent stem cell samples. Biological triplicates of undifferentiated, endoderm-, ectoderm-, and mesoderm-differentiated KOLF2.1J human induced pluripotent stem cells (hiPSCs) were subjected to mass spectrometry. (A) Volcano plot showing differentially abundant proteins between endoderm and ectoderm differentiated hiPSCs. RNAseq- and qPCR-validated endoderm and ectoderm markers are labelled. (B) Volcano plot showing differentially abundant proteins between endoderm and mesoderm differentiated hiPSCs. RNAseq- and qPCR-validated endoderm and mesoderm markers are labelled. (C) Volcano plot showing differentially abundant proteins between ectoderm and mesoderm differentiated hiPSCs. RNAseq- and qPCR-validated ectoderm and mesoderm markers are labelled. (A – C) Proteins are labelled as points in red (significantly less abundant in differentiated vs undifferentiated state), blue (significantly more abundant in differentiated vs undifferentiated state), and grey (no significantly different abundance). Statistical significance was inferred at an “Benjamini Hochberg” post-hoc adjusted p value of ≤ 0.05. n.s.: not significant. logFC: log fold change.

**Supplementary Table 1:**
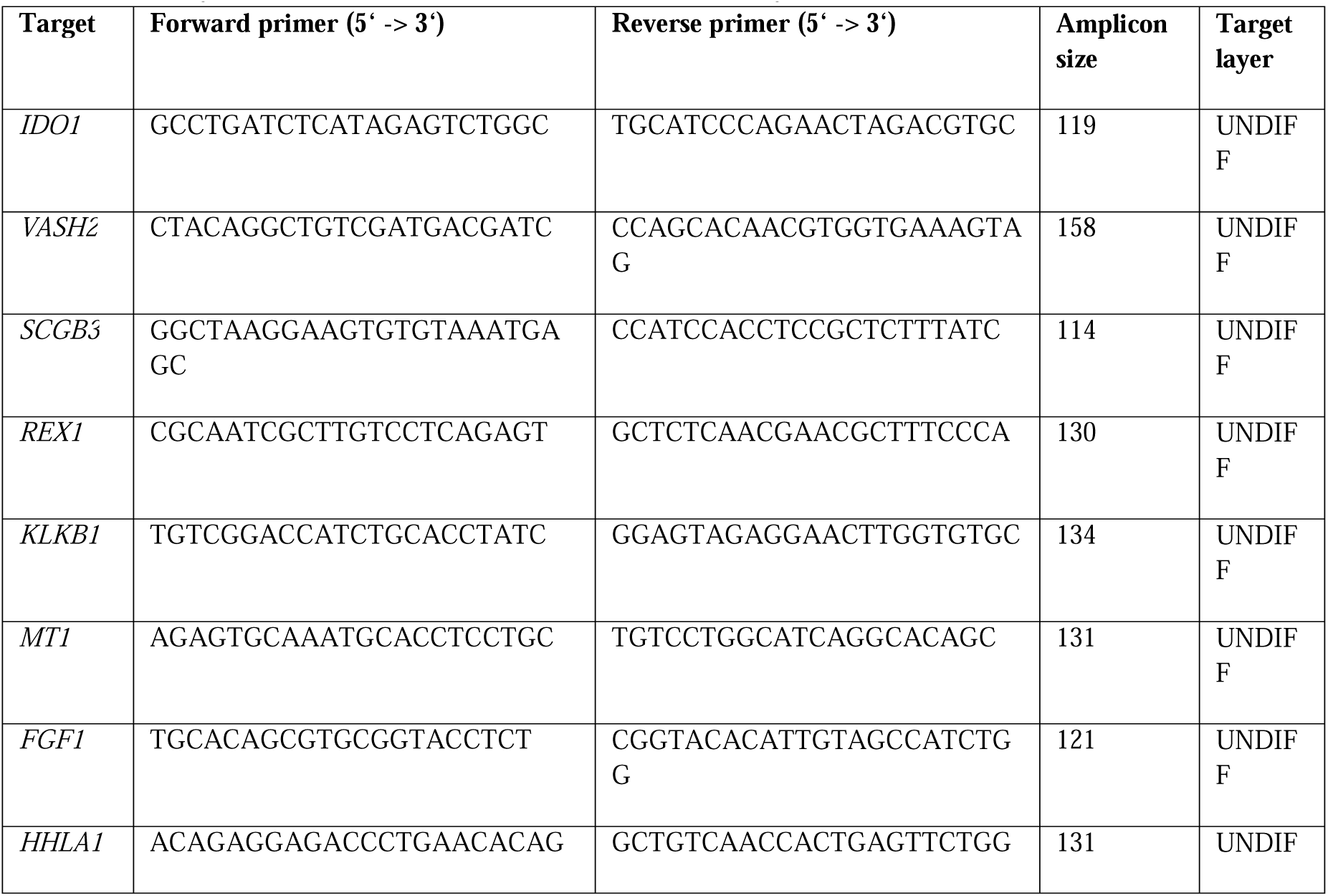
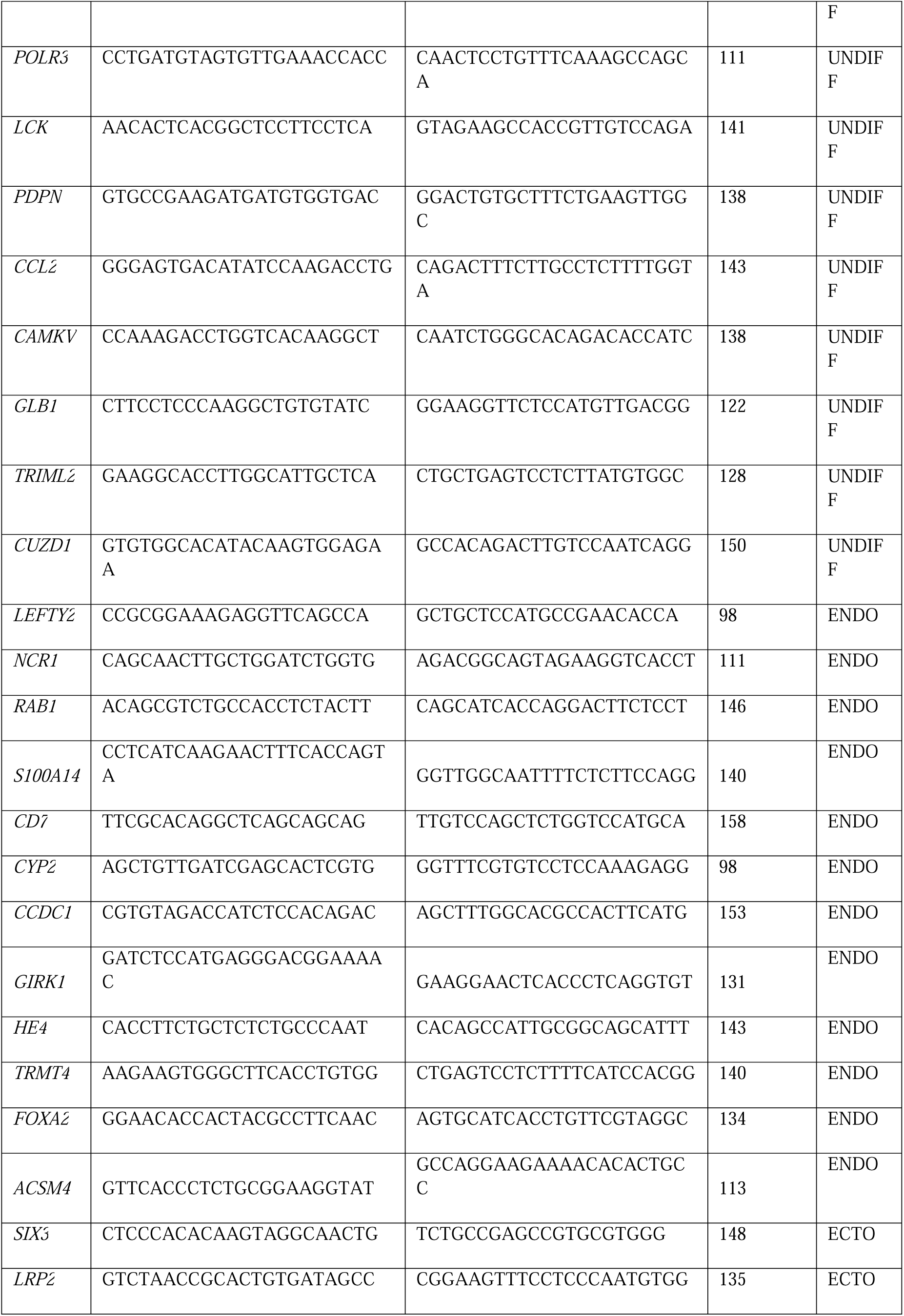
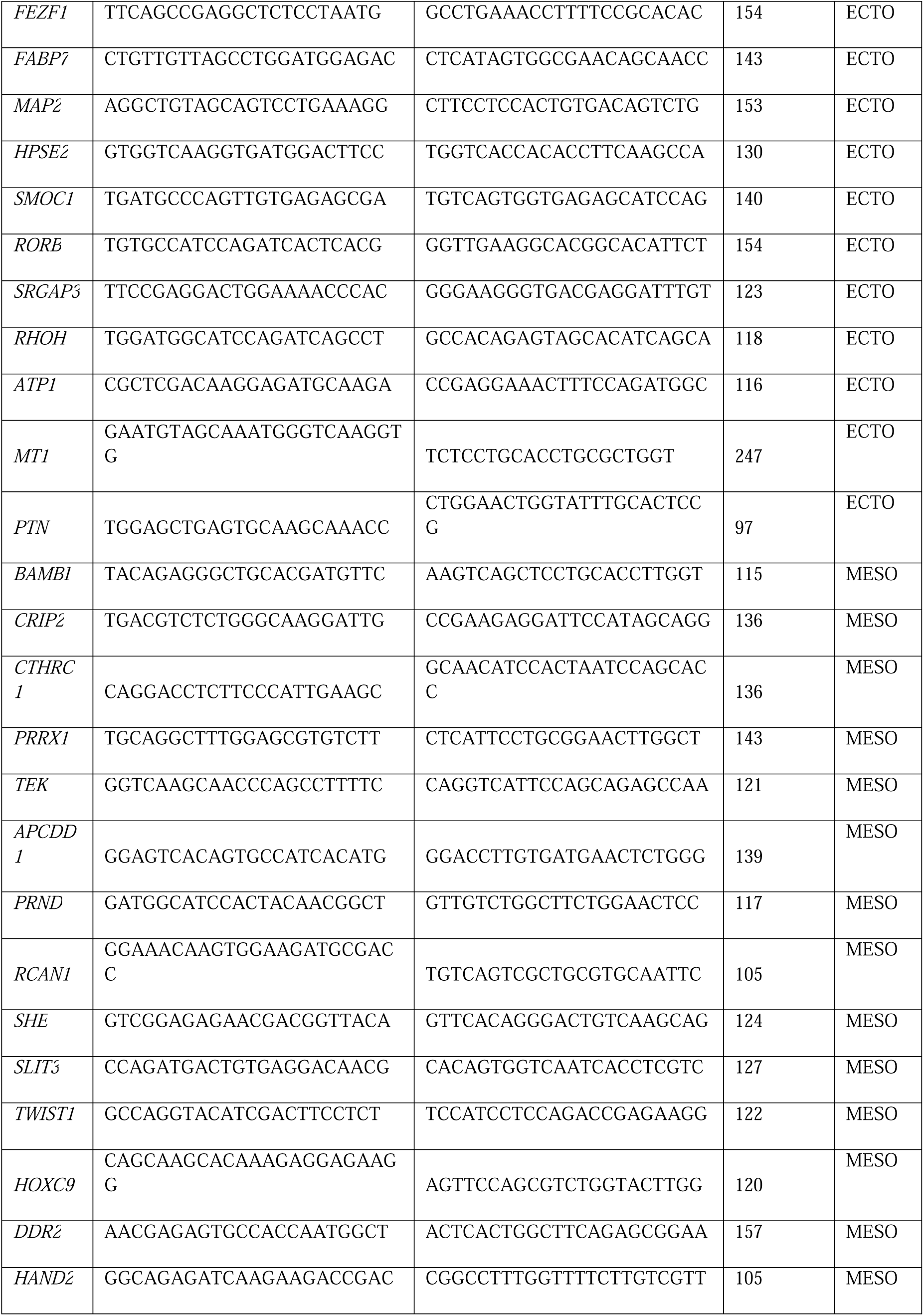
Primer sequences used in this study.

**Supplementary Table 2:**
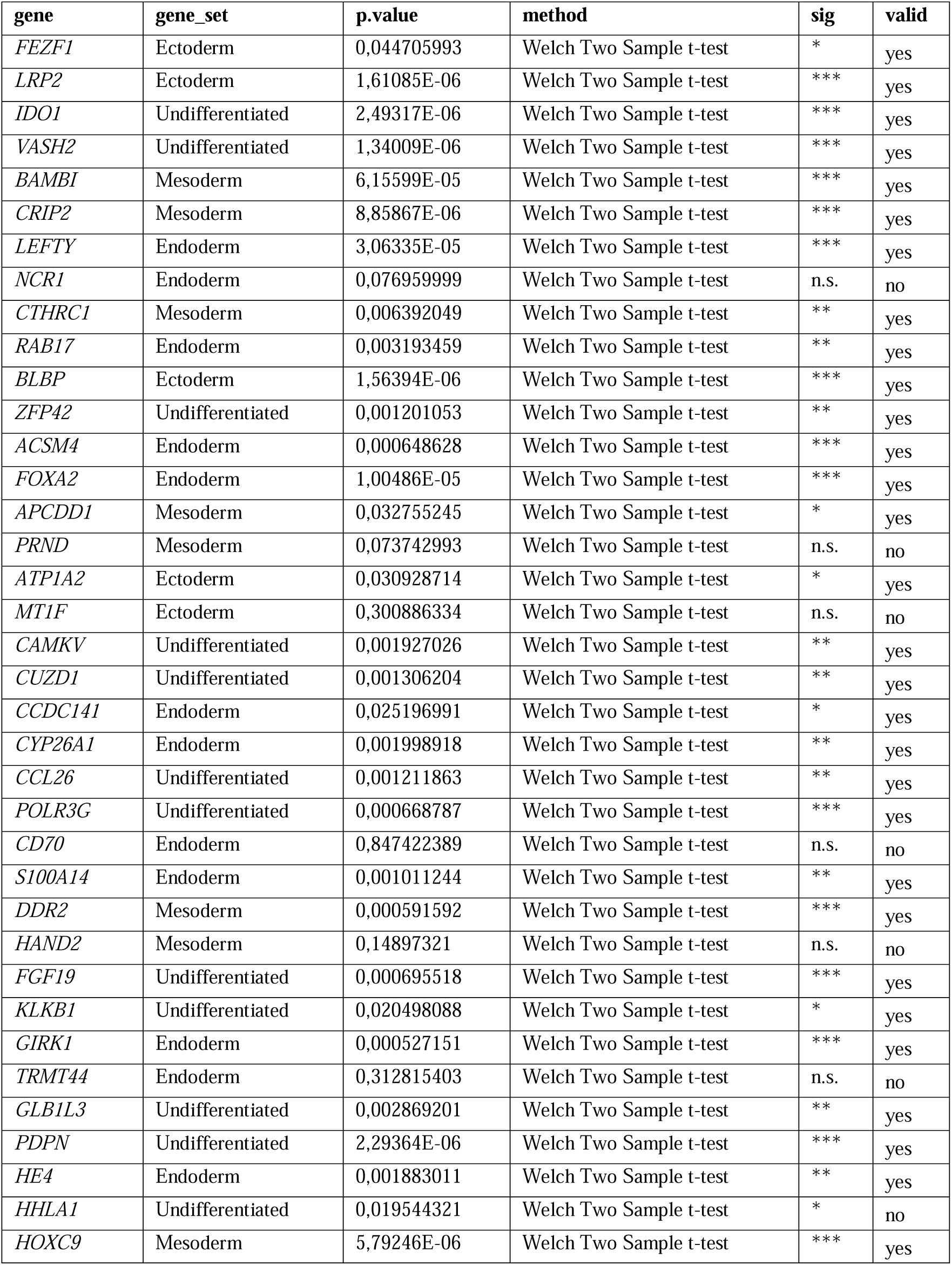
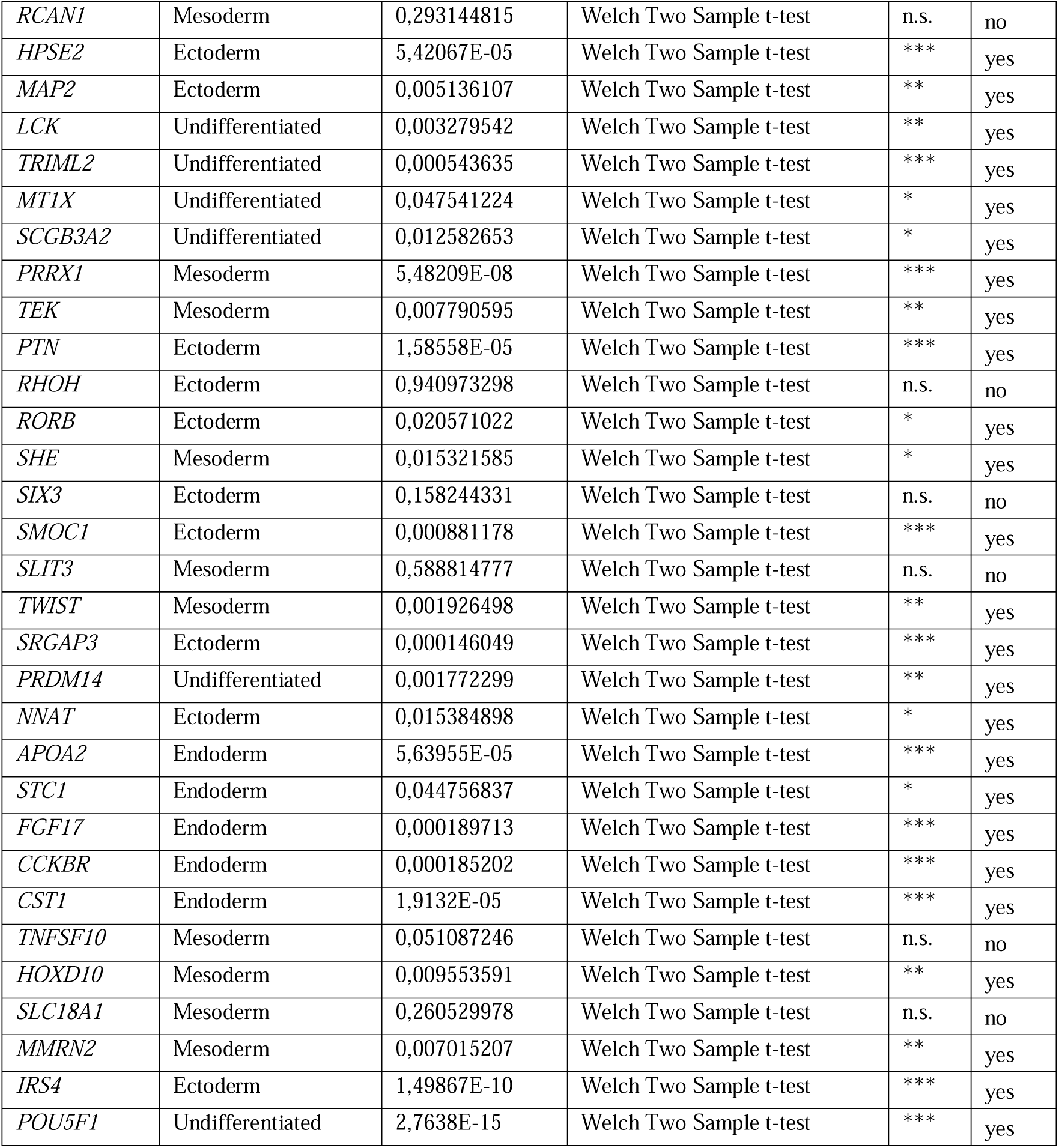
Statistical testing of target genes analyzed in this study.

**Supplementary Table 3: Details on datasets downloaded from gene expression omnibus (GEO).** This table is included as a separate Spreadsheet.

**Supplementary Table 4: Mass Spectrometry measurements for undifferentiated and trilineage-differentiated KOLF2.1J.** This table is included as a separate Spreadsheet.

**Supplementary Table 5:** Cell lines used in this study.

| Cell line | Type | Sex origin | Source | hPSCreg | Reference |
| --- | --- | --- | --- | --- | --- |
| iPSC11 | Commercially available | Male | Cell App. | - | (Ma et al., 2016) |
| iPSC12 | Commercially available | Female | Cell App. | IUFi004-A | (Binder et al., 2024) |
| IMR90 | Commercially available | Female | WiStar | WISCi004-B | - |
| HFF | Reprogrammed from commercially available (Merck) fibroblasts | Male | in-house | - | (Dobner et al., 2024a) |
| DU225 | Genome-edited (parental: iPSC12), *ERCC6*<sup>R683X/R683X</sup> | Female | in-house | - | (Dobner et al., 2024a) |
| DU247 | Genome-edited (parental: iPSC12), KCNK2 KI | Female | in-house | - | (Dobner et al., 2024a) |
| IUFi001 | Reprogrammed from patient fibroblasts | Male | in-house | IUFi001 | (Martins et al., 2021) |
| PGP1 | Commercially available | Male | Synthego | - | (Dobner et al., 2024a) |
| DU22 | Genome-edited (parental: IMR90), Life-Act-GFP | Female | in-house | - | (Tigges, 2021) |
| TFBJ | Reprogrammed from commercially available (ATCC) fibroblasts | Male | in-house | HHUUKDi009-A | (Lorenz et al., 2022) |
| BIHi050 | Reprogrammed from patient fibroblasts | Male | In-house | BIHi050-A | (Dobner et al., 2024a) |
| KOLF2.1J | Commercially available | Male | JAX | WTSli018-B-12 | (Pantazis et al., 2022) |
| AS789 | Reprogrammed from patient fibroblasts (disease: Cockayne syndrome B) | Male | In-house | - | (Szepanowski et al., 2024) |
| SCTi003 | Commercially available | Female | STEMCELL Technologies | SCTi003-A | - |

## Methods

### Human induced pluripotent stem cell culture and differentiation

The human iPSCs analyzed in this study from different sources (Supplementary Table 5) were cultured as described (Dobner et al., 2024a). For maintenance, mTeSR Plus (STEMCELL Technologies) was used as the growth medium, and growth factor-reduced Geltrex (Thermo Fisher) or iMatrix-511 silk (amsbio) were used as matrix coating. All cell lines were routinely tested for mycoplasma contamination by PCR (Minerva Biolabs) or an enzyme-based (Vazyme) method. All cell lines were assessed by G-banding at the start of the study and found to be karyotypically normal. All cell lines were authenticated through short tandem repeat profiling. For directed trilineage differentiation, cells were seeded as described in (Dobner et al., 2024a) and differentiated using commercially available kits (Stemcell Technologies, Miltenyi Biotec).

### RNA isolation, cDNA transcription, and qPCR

After washing with PBS, cells were harvested into Trizol, and the total RNA extracted using a commercially available kit (Zymo). One microgram of total RNA was reverse transcribed into cDNA in a 20 µl reaction (Vazyme), diluted 1:10 with H_2_O, and 1 µl of the dilution used in a 10 µl reaction consisting of 5□µl ChamQ Universal SYBR qPCR Master Mix (Vazyme), 3□µl H_2_O, and 1□µl primer mix (forward□+□reverse at 2.5□µM each for a final primer concentration of 250□nM). Gene expression of target genes was normalized to the mean of two reference genes (*GAPDH*, *ACTB*). Primer sequences and amplicon sizes are listed in Supplementary Table S1.

### Analysis of publicly available bulk and single cell RNAseq datasets

A Gene Expression Omnibus (GEO) database search was conducted using the R package rentrez (version 1.2.4) with the following search term: ‘(germ layer OR ectoderm OR mesoderm OR endoderm) AND “Homo sapiens”[Organism] AND “expression profiling by high throughput sequencing”[DataSet Type] NOT (Chromium OR 10X OR scRNA OR scRNAseq)’. Afterwards, summaries were downloaded and filtered for the terms “hESC|iPSC|pluripotent|ESC” to restrict results to human pluripotent stem cell-derived data. Data was further filtered to identify titles containing “endoderm”, “ectoderm”, and “mesoderm”, and to identify datasets containing either raw or cpm-normalized gene expression data. After this prefiltering, individual datasets were downloaded using the R package GEOquery (version 2.76.0). To assess their suitability for analysis within the presented study based on sample preparation, treatment, and differentiation, supplemental meta data was downloaded using the R package Biobase (version 2.64.0), and manually inspected. If not already cpm-normalized, raw counts were cpm-normalized using the R package edgeR (version 3.40.0). Afterwards, cpm tables were filtered for the validated genes and if no counts were present for a given gene, gene expression values were manually set to 0. Clustering of cpm matrices was performed after log2(+1) transformation and center scaling using the R package ComplexHeatmap (version 2.24.1). A complete list of samples and meta information can be found in Supplementary Table S2.

For the analysis of a gastrulating human embryo, publicly available annotated counts were downloaded from a Carnegie Stage 7 embryo, processed and validated marker genes aggregated to gene expression modules visualized using the R package Seurat (version 5.3.0).

### Flow cytometry of mesoderm-differentiated human induced pluripotent stem cells

For flow cytometry, four different hiPSC lines (SCTi003, iPSC12, KOLF2.1, IMR90) were trilineage-differentiated in duplicates in 24-wells into mesoderm using the StemMACS™ Trilineage Differentiation Kit (Miltenyi Biotec) with the modifications described in (Dobner et al., 2024a). At day 7 of differentiation, cells were washed with PBS, detached using Accutase (PanBiotech), and resuspended in FACS buffer (2% FBS in PBS). After a washing step, cells were distributed into three tubes per cell line per staining condition: single CD144 (FITC-conjugated, Miltenyi Biotec), single CXCR4 (APC-conjugated, Miltenyi Biotec), and double CD144/CXCR4 at 1:50 dilution in FACS buffer for thirty minutes on ice in the dark. After washing twice with cold FACS buffer, cells were analyzed using a FACS Aria III (BD Biosciences). Post-processing and visualization were performed using FlowJo (v 10.10.0).

### Proteome analysis

For mass spectrometry, trilineage differentiation was performed in biological triplicates using the KOLF2.1J reference line in 6-wells according to the manufacturers (Miltenyi) instructions. After the differentiation protocol, at least 1 x 10^6^ cells were prepared and sent for deep mass spectrometry analysis according to the service provider (BGI). In brief, cells were harvested with Accutase, washed with PBS, centrifuged for 10 min at 4 °C at 1,000 G, the supernatant removed and cell pellets snap-frozen in liquid nitrogen for 15 min. Samples were stored at -80 °C and shipped on dry ice. 7,125 proteins were detected on average (range: 6,724 – 7,324). Intensities derived from the service provider (Supplementary Table S3) were subjected to analysis. Missing values were imputed based on (Shi et al., 2025). These values were used for principal component analysis and differential protein level expression using limma (version 3.64.3) by computing empirical Bayes statistics on a linear fit.

### Statistics

For statistical analyses, the open-source script language R was used. For qPCR analysis at least 5 independent biological replicates were analyzed. Differences between groups were calculated using independent t-tests and statistical significance was inferred for p-values ≤ 0.05.

## Resource Availability

Lead contact: Jochen Dobner, Andrea Rossi

## Data and Code Availability

Transcriptome sequencing data was derived from publicly accessible repositories, and the respective sample identifiers are listed in Supplementary Table 3. The DeepDiff web resource can be freely accessed at https://jdobner.shinyapps.io/diff_sets/.

## Acknowledgments

We thank Kira Frye, and Katrin Hollweg for excellent technical assistance.

The IUF is funded by the federal and state governments, the Ministry of Culture and Science of North Rhine-Westphalia (MKW), and the Federal Ministry of Education and Research (BMBF). The Environmental Adaptation and Cellular Resilience lab and the Genome Engineering and Model Development Core Unit are funded by Deutsche Forschungsgemeinschaft (DFG) (RO5380/1-1, RO5380/10-1, RO5380/13-1, RO5380/14-1 to A.R.), AFM-Téléthon (25179 to A.R.), European Union (EU) and North Rhine-Westphalia (NRW) start-up grant (IN-SU-3-001 to A.R and J.D.), VHL (von Hippel-Lindau) betroffener Familien e.V. (to AR) and the Leibniz Competition (SAW) Cooperative Excellence project (K642/2024 to A.R.).

## Author Contributions

J.D.: Conceptualization, Methodology, Software, Validation, Formal analysis, Investigation, Data Curation, Writing – Original Draft, Visualization, Project administration. A.P.: Writing – Review & Editing. A.R.: Conceptualization, Validation, Writing – Original Draft, Supervision, Project administration, Resources, Funding acquisition.

